# [^225^Ac]Ac-SSO110 demonstrates superior efficacy over ^177^Lu-, ^161^Tb-, and ^212^Pb-labeled SSO110 and [^225^Ac]Ac-DOTA-TATE in SSTR2-positive tumor models

**DOI:** 10.64898/2026.03.11.709316

**Authors:** Prachi Desai, Dennis Mewis, Marion Huber, Manuel Sturzbecher-Hoehne, Manfred Ruediger, Germo Gericke, Anika Jaekel

## Abstract

Somatostatin receptor 2 (SSTR2) is highly expressed in neuroendocrine tumors including small cell lung cancer (SCLC) and represents a validated target for peptide receptor radionuclide therapy. The SSTR2 agonist [^177^Lu]Lu-DOTA-TATE is clinically approved, however, treatment resistance and relapse occur. The SSTR2 antagonist SSO110 (DOTA-JR11, OPS201) demonstrates higher tumor uptake and longer retention than DOTA-TATE both pre-clinically and clinically. We performed a systemic head-to-head comparison of SSO110 labeled with various radionuclides of distinct emission characteristics to identify the optimal radionuclide for SSO110 and to compare antagonist with agonist performance.

**Methods:** SSO110 was radiolabeled with ^177^Lu, ^161^Tb, ^212^Pb, and ^225^Ac. Biodistribution was assessed in AR42J and NCI-H69 xenograft models. Therapeutic efficacy of single and fractionated [^212^Pb]Pb-SSO110 was compared with [^177^Lu]Lu-SSO110 in NCI-H69 tumors. Single-dose efficacy of ^225^Ac-, ^161^Tb-, and ^177^Lu-labeled SSO110 was evaluated in both models. [²²⁵Ac]Ac-DOTA-TATE served as agonist comparator. Tumor growth, survival, safety parameters, and tumor absorbed doses were analyzed.

**Results:** All SSO110 radioconjugates demonstrated comparable biodistribution with high tumor uptake and favorable tumor-to-kidney ratios. In NCI-H69 tumors, [^212^Pb]Pb-SSO110 induced dose-dependent tumor growth delay but did not improve anti-tumor efficacy compared with [^177^Lu]u-SSO110 under single or fractionated regimens. [^161^Tb]Tb-SSO110 showed efficacy comparable to [^177^Lu]Lu-SSO110 in NCI-H69 model and significantly improved tumor growth delay in high-SSTR2-expressing AR42J tumors. Across both models, [^225^Ac]Ac-SSO110 demonstrated the highest therapeutic potency, inducing durable tumor regression and 100% survival at clinically relevant activities. [^225^Ac]Ac-SSO110 also outperformed the agonist comparator [^225^Ac]Ac-DOTA-TATE. Dosimetry analysis revealed a 63-fold higher tumor absorbed dose per injected administered activity for [^225^Ac]Ac-SSO110 compared with [^212^Pb]Pb-SSO110. All treatments were well tolerated without significant renal or hepatic toxicity.

**Conclusion:** Therapeutic efficacy of SSTR2-targeted peptide receptor radionuclide therapy appears to benefit from alignment between radionuclide physical half-life and ligand tumor residence time. Among the radionuclides evaluated, [^225^Ac]Ac-SSO110 demonstrated the most pronounced and durable anti-tumor efficacy, outperforming [^161^Tb]Tb-SSO110, [^177^Lu]Lu-SSO110, and the short-lived α-emitter [^212^Pb]Pb-SSO110. These findings support clinical investigation of [^225^Ac]Ac-SSO110 in SSTR2-positive malignancies.

## INTRODUCTION

Somatostatin receptor 2 (SSTR2) is a clinically validated target for imaging and therapy of neuroendocrine tumors (NETs) (*1*,*2*). Beyond gastroenteropancreatic NETs, SSTR2 expression is also observed in several other malignancies, including small-cell lung cancers (SCLC), meningioma, Merkel cell carcinoma, and neuroblastoma, broadening the potential clinical applications of SSTR2-directed radioligand therapy (*3–9*).

SSTR2 agonists such as lutetium-177(¹⁷⁷Lu)-labeled DOTA-TATE (Lutathera^®^) have demonstrated clinical benefit in GEP-NET patients and have been approved as radioligand therapy (RLT) (*10*,*11*). RLT has also been successfully applied to other molecular targets, exemplified by the approval of [^177^Lu]Lu-PSMA-617 (Pluvicto^®^), further supporting the clinical benefit of this therapeutic modality. SSTR2 antagonists such as SSO110 (also known as OPS201 or DOTA-JR11) have emerged as a promising next generation of ligands due to their ability to bind both active and inactive receptor conformations, targeting more binding sites per cell, resulting in higher tumor uptake, and slower dissociation with longer tumor residence time (*12–17*). Clinically, [^177^Lu]Lu-SSO110 has shown markedly higher tumor doses and improved tumor-to-organ ratios compared with [^177^Lu]Lu-DOTA-TATE (*18*).

Several radionuclides are currently being advanced in clinical development for SSTR2-targeted RLT, including β-emitters (^177^Lu), β⁻ emitters with Auger/conversion electrons (terbium-166 (^161^Tb), and α-emitters (actinium-225 (^225^Ac), and lead-212 (^212^Pb)). These payloads vary substantially in linear energy transfer (LET), particle path length, and expected biological effect, providing options to optimally align with the pharmacokinetics of the SSTR2 ligands to maximize tumor cell killing while sparing healthy tissues (*19*,*20*). Comparative studies across SSTR2 antagonist series suggest that ligand-driven pharmacology is largely conserved across radionuclides (*21*,*22*), supporting the systematic evaluation of SSO110 radiolabeled with different therapeutic radionuclides.

^161^Tb with a half-life (t_1/2_) of 6.9 days emits β⁻ particles comparable in energy and half-life to ^177^Lu (t_1/2_: 6.6 days) but additionally releases approximately 11-fold higher numbers of short-range Auger and conversion electrons, enhancing subcellular radiation dose (*23–25*). In contrast, α-emitting radionuclides such as ^225^Ac (t_1/2_: 10.1 days) and ^212^Pb (t_1/2_: 10.6 hours) generate high-LET, micrometer-range (∼50-100 µm) ionization capable of inducing complex and often irreparable DNA damage (*26*).

To determine how different therapeutic radionuclides influence the performance of an SSTR2 antagonist, we evaluated SSO110 radiolabeled with four clinically relevant radionuclides, ^161^Tb, ^177^Lu, ^212^Pb, and ^225^Ac, across two SSTR2-expressing xenograft models that represent distinct biological settings. The NCI-H69 is a SCLC xenograft model characterized by moderate and heterogeneous SSTR2 expression, whereas AR42J is a rat pancreatic xenograft model with high and homogeneous SSTR2 expression. Together, these models enable assessment of SSO110 across the spectrum of SSTR2 density encountered in neuroendocrine malignancies, including SCLC (*4*,*6*). The clinically validated agonist DOTA-TATE was included to contextualize antagonist performance. This framework allows a systematic comparison of therapeutic efficacy for SSO110 across different radionuclide payloads.

## MATERIALS AND METHODS

Detailed information on materials and methods are available in the supplemental data.

### Biodistribution

The biodistribution of radiolabeled SSO110 was evaluated following a single intravenous administration in tumor-bearing nude mice. Swiss nude mice bearing AR42J xenografts received 10 MBq/1 µg ^161^Tb-SSO110, and Balb/c nude mice bearing NCI-H69 xenografts received 450 kBq/1 µg ^212^Pb-, 90 kBq/1 µg ^225^Ac-, or 60 MBq/1 µg ^177^Lu-labeled SSO110.

A constant mass of 1 µg was selected based on prior evidence demonstrating favorable tumor-to-healthy organ uptake ratio for SSO110 at this peptide range (*14*). At predefined time points post-injection, blood, tumor, and selected organs were collected, weighed, and measured for radioactivity. Tissue radioactivity was quantified using a gamma counter for ^161^Tb, ^177^Lu and ^212^Pb, and a liquid scintillation counter for ^225^Ac. Tumor-to-kidney ratios were calculated to evaluate the relative time-course of ^161^Tb-, ^177^Lu-, ^212^Pb-, and ^225^Ac-labeled SSO110. Ex vivo biodistribution results are presented as percentage of injected activity per gram of tissue (%IA/g; mean ± SD).

### Efficacy studies

Mice bearing NCI-H69 or AR42J xenografts received intravenous injections under anesthesia of [^177^Lu]Lu-SSO110, [^161^Tb]Tb-SSO110, [^225^Ac]Ac-SSO110, or [^225^Ac]Ac-DOTA-TATE. The efficacy of single or multiple doses of [^212^Pb]Pb-SSO110 compared with [^177^Lu]Lu-SSO110 was evaluated exclusively in the NCI-H69 xenograft model. Control mice received non-radiolabeled SSO110 or vehicle. An absolute peptide mass of 1 µg was maintained for SSO110 and DOTA-TATE across all treatment groups.

### Tumor absorbed dose calculations

The mean absorbed tumor doses for [^225^Ac]Ac-SSO110 and [^212^Pb]Pb-SSO110 were derived from biodistribution data. Detailed dosimetry calculations are provided in the supplemental data.

## RESULTS

### Biodistribution profile of ^161^Tb-, ^212^Pb-, and ^225^Ac-labeled SSO110 is comparable to [^177^Lu]Lu-SSO110

^212^Pb-, ^225^Ac, ^161^Tb-, and ^177^Lu-labeled SSO110 were synthesized without further purification, yielding radiolabeling incorporation rates of ≥ 95%. To ensure structural consistency with the clinically investigated [^177^Lu]Lu-SSO110 and enable direct comparison of radionuclide-dependent effects without chelator modifications, DOTA was used as the chelator for all radioconjugates.

All biodistribution studies were conducted as independent experiments owing to radionuclide availability and CRO-specific model constraints. Swiss nude mice bearing AR42J xenografts characterized by high and homogenous SSTR2 expression (Fig. S1A) were used to assess [^161^Tb]Tb-SSO110. Balb/c nude mice bearing NCI-H69 xenografts with moderate and heterogenous SSTR2 expression (Fig. S1B) were used to assess ^225^Ac-, ^212^Pb, and ^177^Lu-labeled SSO110. Accordingly, biodistribution data are interpreted within each model context.

Following intravenous injection, all radioconjugates rapidly cleared from blood with < 0.3 %IA/g remaining at 4 h (Fig. 1). High tumor uptake was observed for all compounds, with peak uptake at 1 h for [^212^Pb]Pb-SSO110 (6.0 %IA/g) and at 4 h post injection for ^225^Ac-, ^161^Tb-, and ^177^Lu-labeled SSO110 (10.3, 18.3, and 5.7 %IA/g, respectively). The higher tumor uptake observed for [^161^Tb]Tb-SSO110 in the AR42J model is consistent with the higher SSTR2 expression in this xenograft. Supporting this interpretation, internal biodistribution studies with [^177^Lu]Lu-SSO110 have consistently demonstrated higher tumor uptake in AR42J compared with NCI-H69 tumors (data not shown).

As a consequence of urinary excretion and partial tubular reabsorption, high renal uptake was observed, with peak values at 1 h for [^212^Pb]Pb-SSO110 (14.7 %IA/g) and at 4 h for ^225^Ac-, ^161^Tb-, and ^177^Lu-labeled SSO110 (13.9, 9.6, and 8.8 %IA/g, respectively). Moderate uptake (< 5 %IA/g) was detected in SSTR2-expressing organs such as pancreas and stomach across all radioconjugates (Fig. 1). Radioactivity in all organs rapidly declined over time. Tumor-to-kidney ratios increased over time for [^161^Tb]Tb-SSO110, [^177^Lu]Lu-SSO110, and [^225^Ac]Ac-SSO110, whereas [^212^Pb]Pb-SSO110 did not show a relative increase (Figure S2).

Overall, these findings indicate that SSO110 maintains a consistent biodistribution profile across all radionuclides tested.

**Figure 1.**
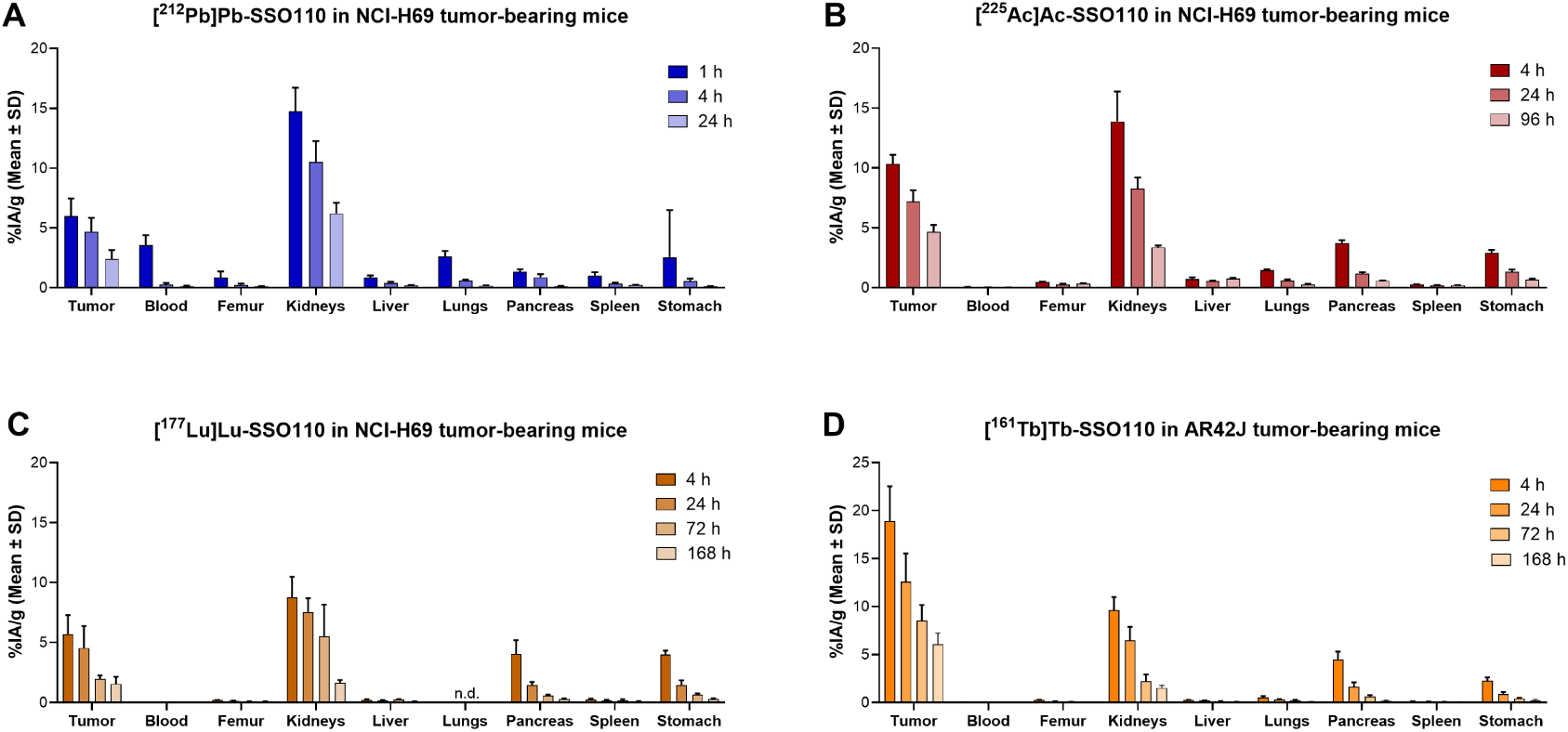
Biodistribution of ^212^Pb-, ^225^Ac-, ^177^Lu-, and ^161^Tb-labeled SSO110. %IA/g of selected organs after administration of (A) 450 kBq [^212^Pb]Pb-SSO110 (n=5), (B) 90 kBq [^225^Ac]Ac-SSO110 (n=4), (C) 60 MBq [^177^Lu]Lu-SSO110 (n=4), and (D) 10 MBq [^161^Tb]Tb-SSO110 (n=4). Values are represented as mean ± SD.

### [^212^Pb]Pb-SSO110 achieves anti-tumor activity comparable to [^177^Lu]Lu-SSO110

The efficacy of single escalating doses of [^212^Pb]Pb-SSO110 (0.1, 0.24, and 0.5 MBq) and a single dose of [^177^Lu]Lu-SSO110 (21.5 MBq) was evaluated in the NCI-H69 xenograft model with moderate and heterogenous SSTR2 expression. The selected [^212^Pb]Pb-SSO110 activity range was based on previously reported efficacious doses of ^212^Pb-labeled somatostatin analogs in preclinical models and was designed to assess dose-dependent effects within a tolerable activity window (*27*). An administered activity of 21.5 MBq [^177^Lu]Lu-SSO110 was selected to reflect a clinically relevant dose, corresponding to approximately 4.5 GBq used in ongoing clinical trials. Treatment with [^212^Pb]Pb-SSO110 induced a dose-dependent delay in tumor growth. At the highest activity tested, 0.5 MBq [^212^Pb]Pb-SSO110 achieved tumor volume reduction and median survival (50 days) comparable to 21.5 MBq [^177^Lu]Lu-SSO110 (47 days), indicating similar single-dose efficacy of the two radioconjugates in prolonging survival at the doses tested (Fig. 2A, 2B).

**Figure 2.**
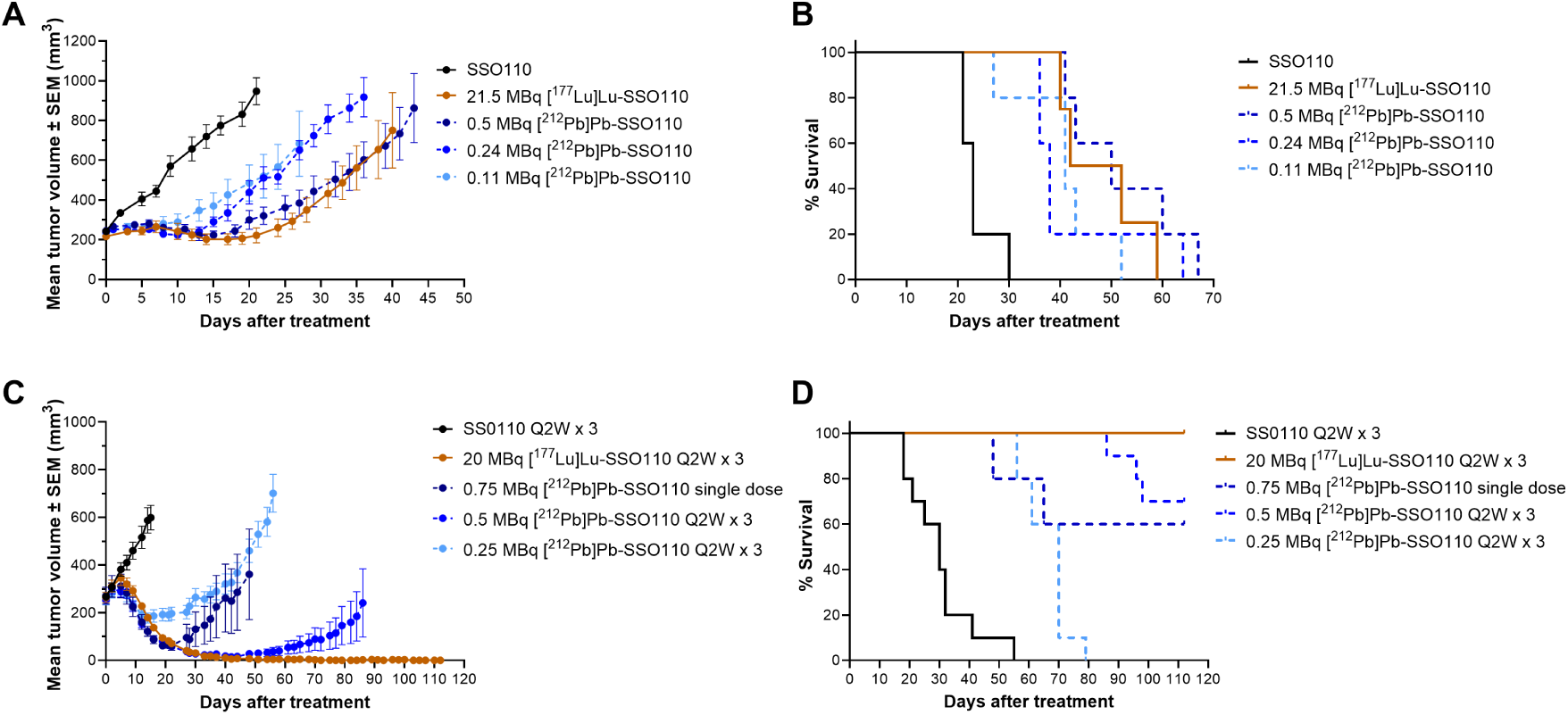
Tumor growth and survival following treatments in the NCI-H69 xenograft model. (A) Tumor volumes and (B) Kaplan-Meier survival analysis of NCI-H69 xenograft-bearing mice treated with a single-dose of 21.5 MBq [^177^Lu]Lu-SSO110 or 0.1 - 0.5 MBq [^212^Pb]Pb-SSO110. (C) Tumor volumes and (D) Kaplan-Meier survival analysis of NCI-H69 xenograft-bearing mice treated with fractionated dosing (Q2W x 3) of 20 MBq [^177^Lu]Lu-SSO110 or 0.25 and 0.5 MBq [^212^Pb]Pb-SSO110, or with single dose of 0.75 MBq [^212^Pb]Pb-SSO110. Tumor volumes are shown as mean ± SEM (n=5-6 for A and C; n=10 for B and D) and are plotted until the first animal dropout.

In addition, the impact of fractionated dosing was assessed using [^212^Pb]Pb-SSO110 (0.25 and 0.5 MBq) and [^177^Lu]Lu-SSO110 (20 MBq) administered every two weeks for a total of three doses (Q2W x 3) and compared with a single dose of 0.75 MBq [^212^Pb]Pb-SSO110. Control animals received three injections of non-radiolabeled SSO110. Fractionated treatment with 20 MBq [^177^Lu]Lu-SSO110 (Q2W x 3) induced complete tumor remissions in 100% of mice, whereas 0.5 MBq [^212^Pb]Pb-SSO110 (Q2W x 3) achieved complete responses in 50% of animals, with tumor regrowth observed in the remaining mice. A single 0.75 MBq dose of [^212^Pb]Pb-SSO110 induced an initial tumor volume reduction that was more pronounced than multiple doses of 0.25 MBq [^212^Pb]Pb-SSO110, but less effective than fractionated doses of 20 MBq [^177^Lu]Lu-SSO110 (Fig. 2C,D). All treatments were well tolerated, with no major body-weight loss observed (Fig. S3).

### [^225^Ac]Ac-SSO110 demonstrates durable complete remissions compared to ^177^Lu- and ^161^Tb-labeled SSO110 and [^225^Ac]Ac-DOTA-TATE in the NCI-H69 xenograft model

The therapeutic efficacy of SSO110 labeled with the β-emitters ^161^Tb and ^177^Lu and the α-emitter ^225^Ac was evaluated in mice bearing NCI-H69 xenografts following single-dose administration. To enable direct comparison of ß-emitting radionuclides, [^161^Tb]Tb-SSO110 and [^177^Lu]Lu-SSO110 were administered at identical activities of 20 MBq. For α-therapy evaluation, [^225^Ac]Ac-SSO110 was tested at 21 and 42 kBq. The 42 kBq dose corresponds to a clinically relevant human-equivalent activity of approximately 10.2 MBq. The SSTR2 agonist [^225^Ac]Ac-DOTA-TATE (42 kBq) was included as a clinically established comparator.

Vehicle-treated mice showed progressive tumor growth and reached a median survival of 36 days. All radiolabeled SSO110 constructs induced anti-tumor responses (Fig. 3). Notably, therapeutic outcome was strongly radionuclide-dependent.

The β-emitting constructs [^161^Tb]Tb-SSO110 and [^177^Lu]Lu-SSO110 demonstrated comparable tumor-growth delay and median survival, indicating similar therapeutic performance in this model. In contrast, α-emitting [^225^Ac]Ac-SSO110 induced complete tumor regressions at 42 kBq resulting in 100% survival, with 50% of mice remaining tumor-free at study termination (day 70). Even at 21 kBq, substantial anti-tumor activity was achieved, underscoring the high therapeutic potency of [^225^Ac]Ac-SSO110.

Importantly, when directly compared at identical activity levels, [^225^Ac]Ac-SSO110 demonstrated superior efficacy relative to [^225^Ac]Ac-DOTA-TATE. While [^225^Ac]Ac-DOTA-TATE, [^177^Lu]Lu-SSO110, and [^161^Tb]Tb-SSO110 showed broadly comparable tumor-growth delay and survival benefit at the tested doses, only [^225^Ac]Ac-SSO110 achieved durable complete remissions.

All treatments were well tolerated, with no treatment-related adverse effects observed relative to vehicle. Body weights increased throughout the study, indicating good tolerability (Fig S4A). Clinical chemistry analysis including ALB, ALP, BUN, CREA, and TBIL revealed no relevant differences between treatment groups and vehicle controls (Table 1).

**Figure 3.**
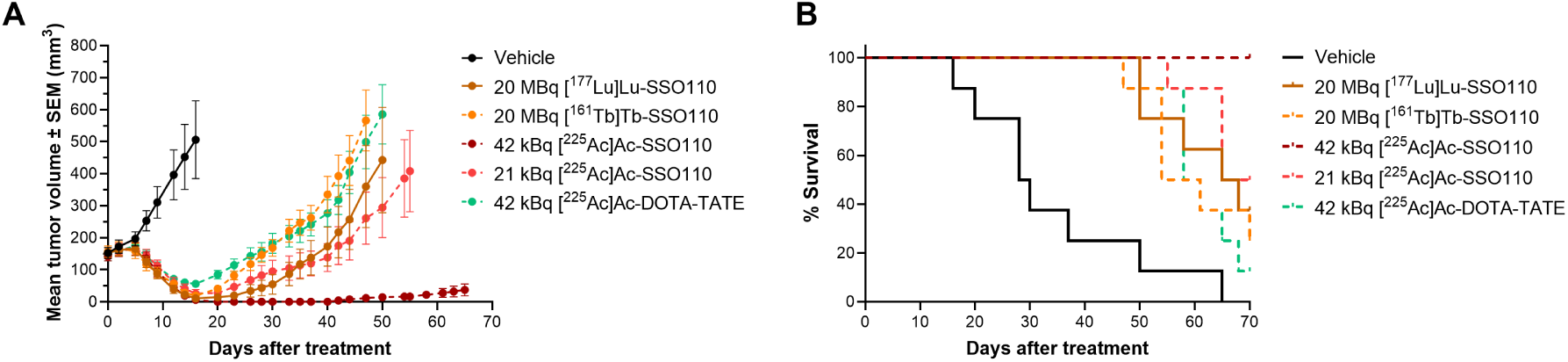
Tumor growth and survival following treatment in the NCI-H69 xenograft model. Tumor volumes and (B) Kaplan-Meier survival curves of NCI-H69 xenograft-bearing mice treated with vehicle, 20 MBq ^177^Lu- or ^161^Tb-labeled SSO110, 21 and 42 kBq [^225^Ac]Ac-SSO110, and 42 kBq [^225^Ac]Ac-DOTA-TATE. Tumor volumes are shown as mean ± SEM (n=8) and plotted until the first animal dropout.

**Table 1.** Plasma clinical chemistry parameters in NCI-H69 xenograft-bearing mice administered with vehicle or radioconjugates (mean ± SD).

| Treatment groups | ALB<br>(g/L) | ALP<br>(U/L) | BUN<br>(mmol/L) | CREA<br>(μmol/L) | TBIL<br>(μmol/L) |
| --- | --- | --- | --- | --- | --- |
| Vehicle | 27.63 ± 0.86 | 58.75 ± 6.20 | 7.64 ± 1.33 | 18.06 ± 8.80 | 1.06 ± 1.24 |
| 20 MBq [<sup>177</sup>Lu]Lu-SSO110 | 27.31 ± 0.77 | 61.25 ± 5.06 | 6.85 ± 0.83 | 23.70 ± 5.13 | 0.50 ± 0.60 |
| 20 MBq [<sup>161</sup>Tb]Tb-SSO110 | 28.66 ± 1.84 | 56.88 ± 4.22 | 7.16 ± 1.03 | 14.54 ± 3.43 | 1.47 ± 1.07 |
| 21 kBq [<sup>225</sup>Ac]Ac-SSO110 | 27.71 ± 1.29 | 62.38 ± 6.52 | 7.33 ± 0.87 | 24.23 ± 8.61 | 0.79 ± 0.71 |
| 42 kBq [<sup>225</sup>Ac]Ac-SSO110 | 26.21 ± 1.49 | 67.00 ± 12.18 | 7.17 ± 1.74 | 18.79 ± 8.64 | 1.51 ± 1.65 |
| 42 kBq [<sup>225</sup>Ac]Ac-DOTA-TATE | 28.00 ± 1.36 | 54.63 ± 7.94 | 7.63 ± 0.89 | 12.85 ± 5.98 | 0.90 ± 0.75 |

### Superior anti-tumor efficacy of [^225^Ac]Ac-SSO110 in the AR42J model compared with other radioconjugates

To evaluate the impact of radionuclide selection under conditions of high target expression, single escalating doses of [^225^Ac]Ac-SSO110 (10, 21, and 37 kBq) were assessed in the SSTR2^high^AR42J xenograft model. For radionuclide comparison, single doses of [^161^Tb]Tb-SSO110 and [^177^Lu]Lu-SSO110 at 20 MBq each were administered, and [^225^Ac]Ac-DOTA-TATE at 37 kBq was included as benchmark.

Vehicle-treated animals showed rapid tumor progression with a median survival of 11 days. In contrast to the NCI-H69 xenograft model, radionuclide-dependent differences among beta-emitters became apparent in AR42J tumors. At identical activity levels, 20 MBq [^161^Tb]Tb-SSO110 significantly prolonged tumor growth delay and median survival compared to 20 MBq [^177^Lu]Lu-SSO110 (36 vs. 30.5 days, p=0.0194) (Fig. 4A, B), indicating enhanced efficacy of ^161^Tb in the high-SSTR2 setting.

Treatment with [^225^Ac]Ac-SSO110 resulted in dose-dependent anti-tumor responses. A single dose as low as 10 kBq [^225^Ac]Ac-SSO110 induced tumor growth delay comparable to 20 MBq [^177^Lu]Lu-SSO110. At activities above 10 kBq, [^225^Ac]Ac-SSO110 exceeded the efficacy of [^161^Tb]Tb-SSO110.

Notably, 37 kBq [^225^Ac]Ac-SSO110 induced immediate tumor growth arrest that persisted until study termination (48 days), resulting in 100% survival. In contrast, tumors treated with 37 kBq [^225^Ac]Ac-DOTA-TATE resumed growth by day 20, resulting in a median survival of 33 days. Importantly, [^225^Ac]Ac-SSO110 demonstrated superior efficacy relative to [^225^Ac]Ac-DOTA-TATE, even at 50% lower administered activity. At the doses tested, 20 MBq [^177^Lu]Lu-SSO110 and 20 MBq [^161^Tb]Tb-SSO110 showed anti-tumor efficacy and prolongation of median survival comparable to 37 kBq [^225^Ac]Ac-DOTA-TATE, whereas only [^225^Ac]Ac-SSO110 achieved sustained tumor-growth suppression.

Body weights increased throughout treatment, indicating good tolerability (Fig. S4B). No treatment-related increases were observed in serum clinical chemistry parameters (ALB, ALP, BUN, CREA or TBIL). A modest decrease in ALP (17% compared to vehicle) was noted in the 20 MBq [^161^Tb]Tb-SSO110 and [^225^Ac]Ac-DOTA-TATE groups (Table 2); however, this change was not accompanied by alterations in other hepatic or renal parameters and correlated with the absence of treatment-related findings in liver and kidney histopathology (data not shown).

Histopathological analysis of AR42J tumors collected at the day of euthanasia showed increased necrosis (Fig. S5), reduced mitotic index, and a higher proportion of regressing neoplastic cell phenotype following treatment with 37 kBq [^225^Ac]Ac-SSO110 compared to 37 kBq [^225^Ac]Ac-DOTA-TATE and vehicle (Fig. 5A,B).

**Figure 4.**
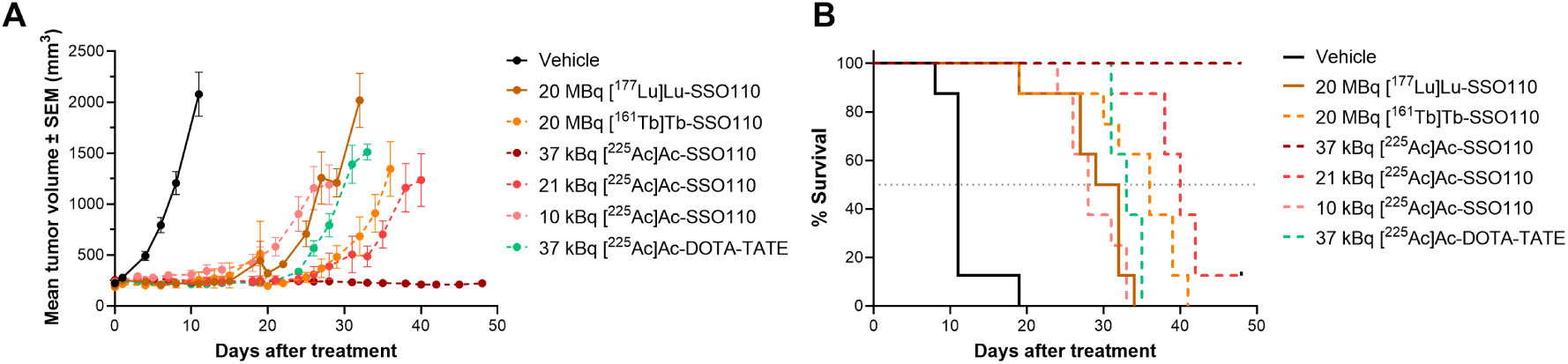
Tumor growth and survival following treatment in the AR42J xenograft model. (A) Tumor volumes and (B) Kaplan-Meier survival curves of AR42J xenograft-bearing mice treated with 20 MBq of ^161^Tb- or ^177^Lu-labeled SSO110, 10, 21, and 37 kBq [^225^Ac]Ac-SSO110, and 37 kBq [^225^Ac]Ac-DOTA-TATE. Tumor volumes are presented as mean ± SEM (n=7-8) and are plotted until 50% of animals remained in the study.

**Table 2.**
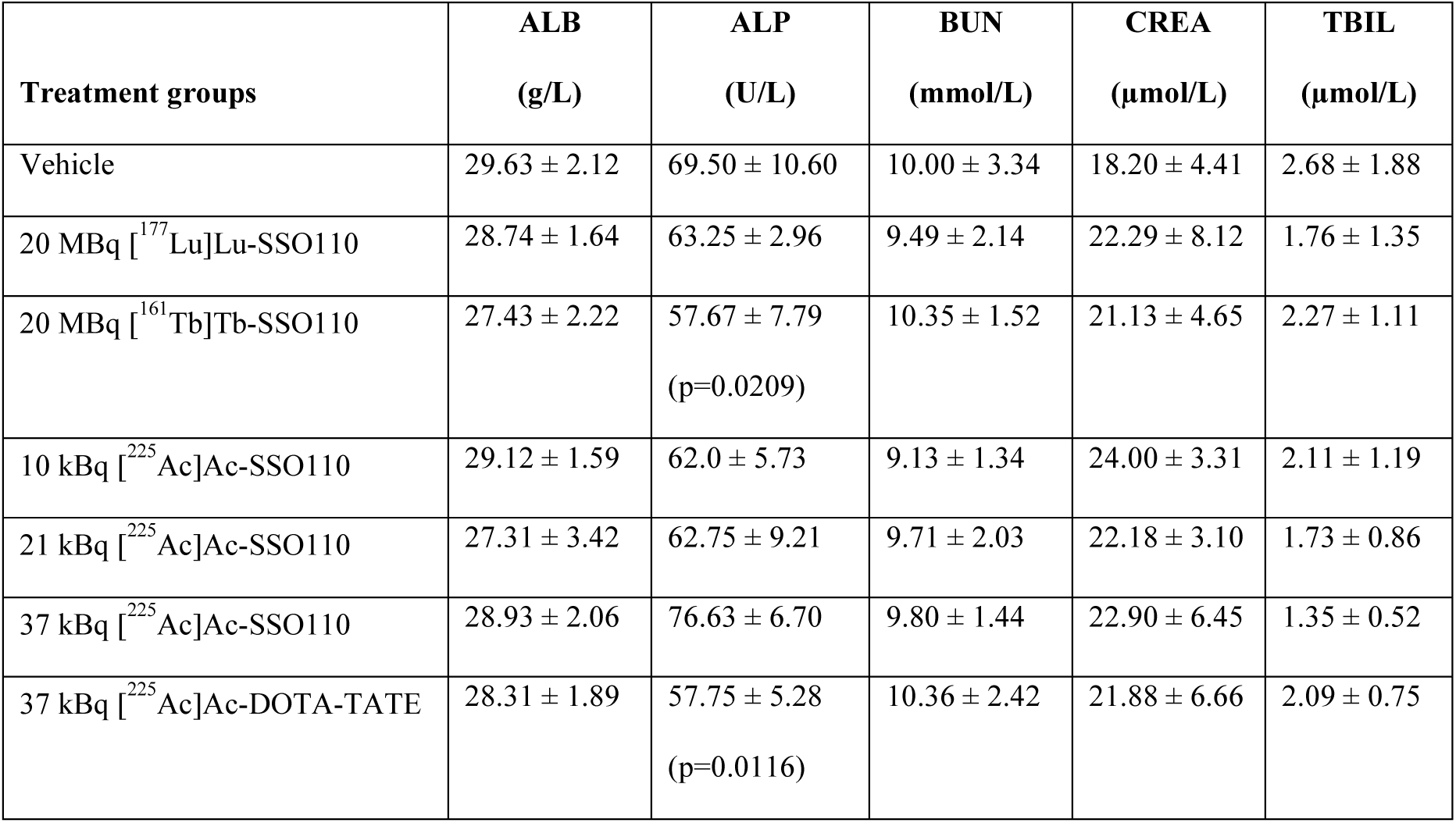
Serum clinical chemistry parameters of AR42J xenograft-bearing mice administered with vehicle or radioconjugates (mean ± SD).

**Figure 5.**
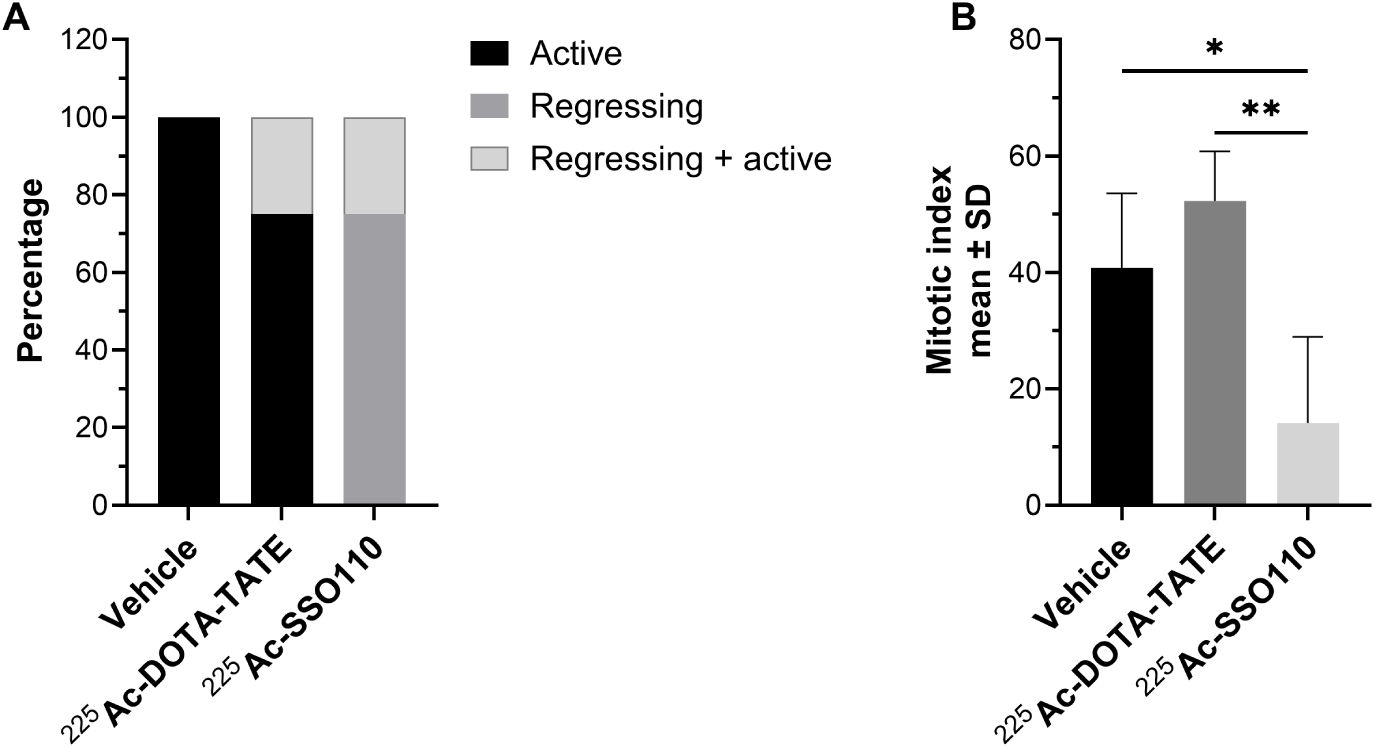
Histopathological analysis of AR42J xenograft tumor sections after treatment with vehicle, 37 kBq [^225^Ac]Ac-DOTA-TATE or [^225^Ac]Ac-SSO110. (A) Neoplastic cell phenotype expressed as the percentage of active, regressing or active plus regressing cells in tumor sections. (B) Mitotic index of neoplastic cells. Data are presented as mean ± SD.

### [^225^Ac]Ac-SSO110 delivers higher tumor absorbed doses than [^212^Pb]Pb-SSO110

To mechanistically explain the superior therapeutic efficacy of [^225^Ac]Ac-SSO110 relative to [^212^Pb]Pb-SSO110, despite both being α-emitting radioconjugates, tumor absorbed dose estimates were derived from biodistribution-based tumor time-activity curves in NCI-H69 xenografts using the MIRD formalism. Based on these analyses, [^225^Ac]Ac-SSO110 delivered a 63-fold higher mean tumor absorbed dose per injected kBq than [^212^Pb]Pb-SSO110 (Table 3). These dosimetry findings provide a quantitative rationale for the markedly superior anti-tumor efficacy of [^225^Ac]Ac-SSO110 observed in this study and emphasize the importance of radionuclide-ligand pharmacokinetic compatibility for effective therapy.

**Table 3.** Mean tumor absorbed dose per injected kBq (mean ± SD)

| Radioconjugates | Mean tumor absorbed dose (Gy*kBq<sup>-1</sup>) |
| --- | --- |
| [<sup>225</sup>Ac]Ac-SSO110 | 1.62*10<sup>-1</sup> ± 1.20*10<sup>-2</sup> |
| [<sup>212</sup>Pb]Pb-SSO110 | 2.57*10<sup>-3</sup> ± 1.75*10<sup>-4</sup> |

## DISCUSSION

In this study, we performed a systematic head-to-head comparison of the SSTR2 antagonist SSO110 labeled with radionuclides of distinct emission profiles, including the β-emitter ^177^Lu, the β- and Auger/conversion electron emitter ^161^Tb, and the α-emitters ^212^Pb and ^225^Ac, across two SSTR2-positive xenograft models. In parallel, the clinically established agonist DOTA-TATE was included as a benchmark. Our data demonstrate that radionuclide selection profoundly influences therapeutic efficacy of SSO110 and that [^225^Ac]Ac-SSO110 exhibits the most pronounced and durable anti-tumor activity across models.

Importantly, radiolabeling with different therapeutic radionuclides did not alter the biodistribution of SSO110, as ^161^Tb-, ^212^Pb-, and ^225^Ac-labeled constructs showed profiles highly comparable to that of [^177^Lu]Lu-SSO110. Radiolabeled SSO110 accumulated efficiently in SSTR2-expressing xenograft tumors, and in kidneys, consistent with the known renal clearance pathway of SSO110 and other peptide-based radiopharmaceuticals. Uptake was also observed in SSTR2-expressing organs such as pancreas and stomach, whereas little to no uptake was observed in the liver, lungs, or other non-target organs. Notably, ^161^Tb-, ^177^Lu-, and ^225^Ac-labeled SSO110 cleared more rapidly from the kidneys than from tumor tissue, resulting in increasing tumor-to-kidney ratios over time. In contrast, tumor-to-kidney ratios for [^212^Pb]Pb-SSO110 did not improve over time, likely due to the short physical half-life of ^212^Pb (10.6 h) relative to the long tumor retention of SSO110. Together, these findings indicate that radiolabeling with different therapeutic radionuclides does not alter the in vivo targeting properties of SSO110. Similar observations were reported by Handula et al. in a direct comparison of ^225^Ac- and ^177^Lu-labeled DOTA-JR11 (*28*).

Despite its α-emission characteristics, [^212^Pb]Pb-SSO110 did not improve tumor growth delay or survival relative to [^177^Lu]Lu-SSO110 in the NCI-H69 model. Although administration of [^212^Pb]Pb-SSO110 induced dose-dependent tumor growth inhibition, the activity required to achieve anti-tumor efficacy comparable to or greater than [^177^Lu]Lu-SSO110 in the NCI-H69 xenograft model (0.75 MBq [^212^Pb]Pb-SSO110 per 20 g mouse) corresponds to an estimated human-equivalent activity of approximately 183 MBq for a 60 kg patient, based on body surface area conversion as recommended in FDA guidance for dose conversion of molecules below 100,000 Da. This exceeds clinically investigated activity levels for ^212^Pb-based therapies such as those evaluated in the ALPHAMEDIX02 clinical trial (NCT05153772) at 2.5 MBq/kg, corresponding to approximately 150 MBq for a 60 kg patient. While further dose escalation might enhance therapeutic efficacy of [^212^Pb]Pb-SSO110, such activity levels may not be feasible preclinically and clinically due to anticipated safety and tolerability constraints. These results suggest that the limited efficacy of [^212^Pb]Pb-SSO110 is attributed to the kinetic mismatch between its short physical half-life and the prolonged tumor retention of SSO110, limiting the number of effective radioactive decays occurring within the tumor.

In contrast, [^225^Ac]Ac-SSO110 demonstrates the strongest therapeutic efficacy in comparison with ^212^Pb-, ^177^Lu-, and ^161^Tb-labeled SSO110. [^225^Ac]Ac-SSO110 induced dose-dependent anti-tumor responses, with marked tumor growth inhibition observed at activities as low as 10 kBq (human equivalent dose of 2.4 MBq per 60 kg patient). Durable tumor regressions were achieved at clinically relevant activity levels in both NCI-H69 and AR42J xenograft models, resulting in 100% overall survival. Importantly, [^225^Ac]Ac-SSO110 was well tolerated, with no clinical signs of acute toxicity and no evidence of liver or kidney damage based on clinical chemistry and histopathological evaluation, supporting a favorable therapeutic window.

To better understand the observed differences in therapeutic potency between ^212^Pb- and ^225^Ac-labeled SSO110, tumor uptake data from the biodistribution studies were used to estimate the tumor absorbed doses. [^225^Ac]Ac-SSO110 delivered a 63-fold higher tumor absorbed dose per administered kBq compared with [^212^Pb]Pb-SSO110. This difference far exceeds the approximately 3.5-fold higher emitted energy per decay of ^225^Ac relative to ^212^Pb, indicating that the superior efficacy of [^225^Ac]Ac-SSO110 is primarily driven by a greater number of radioactive decays occurring within the tumor.

Comparison of therapeutic potency between ^161^Tb- and ^177^Lu-labeled SSO110 revealed model-dependent differences. In the NCI-H69 xenograft model, both radioconjugates showed comparable anti-tumor activity and survival benefit. In contrast, in the AR42J xenograft model, [^161^Tb]Tb-SSO110 significantly prolonged tumor growth delay and median survival compared with [^177^Lu]Lu-SSO110, consistent with previous findings using the SSTR2 antagonists LM3 (*25*). This model-specific difference could be attributed to a more homogeneous SSTR2 expression in AR42J tumors, potentially increasing the therapeutic relevance of the short-range Auger and conversion electrons emitted by ^161^Tb. Nevertheless, even in the AR42J model, [^225^Ac]Ac-SSO110 demonstrated superior efficacy compared with [^161^Tb]Tb-SSO110, highlighting the dominant contribution of sustained α-particle dose delivery when pharmacokinetically aligned with tumor retention.

Direct comparison of [^225^Ac]Ac-SSO110 with [^225^Ac]Ac-DOTA-TATE demonstrated that α-emission alone does not explain therapeutic superiority. In both xenograft models, [^225^Ac]Ac-SSO110 induced durable tumor regressions and improved survival compared with [^225^Ac]Ac-DOTA-TATE. Complete tumor remissions and 100% overall survival were observed only with [^225^Ac]Ac-SSO110, whereas tumors treated with [^225^Ac]Ac-DOTA-TATE resumed growth and showed shorter median survival. Enhanced efficacy of SSO110 relative to DOTA-TATE has also been observed preclinically for β-emitters (*13*,*14*), indicating that the superiority of SSO110 to DOTA-TATE is independent of radionuclide choice. The improved performance of SSO110 can be explained by its antagonist binding profile. SSO110 binds to both active and inactive SSTR2 conformations, resulting in an approximately four-fold higher number of accessible binding sites per cell and increased receptor occupancy, as well as a markedly slower dissociation rate compared with DOTA-TATE (*16*). These properties translate into enhanced tumor uptake and prolonged tumor retention compared with the agonist DOTA-TATE, as observed preclinically and clinically (*13*,*14*,*18*). This is further supported by clinical observations from a phase I/II study of [^177^Lu]Lu-SSO110, in which therapeutic effects comparable to Lutathera^®^ were achieved despite administration of lower activity of [^177^Lu]Lu-SSO110 (*29*). Collectively, these properties of SSO110 enable higher tumor dose delivery compared to DOTA-TATE despite minimal internalization, challenging the long-standing paradigm that receptor internalization is required for therapeutic efficacy of radiopharmaceuticals. Such prolonged tumor residence may be particularly relevant for α-emitters such as ^225^Ac, as extended retention increases the fraction of decays occurring within the tumor microenvironment. Overall, these findings indicate that therapeutic performance of radioconjugates is governed by the alignment of radionuclide emission characteristics and half-life with ligand-mediated tumor retention.

## CONCLUSION

This comprehensive preclinical comparison demonstrates that radionuclide selection and ligand-specific tumor retention jointly determine therapeutic efficacy of SSTR2-targeted radiopharmaceutical therapy. Although biodistribution profiles were comparable across radionuclides, therapeutic outcome differed markedly. Among all tested radioconjugates, [^225^Ac]Ac-SSO110 exhibited the most pronounced and durable anti-tumor efficacy, outperforming ^212^Pb-, ^161^Tb- and ^177^Lu-labeled SSO110 as well as [^225^Ac]Ac-DOTA-TATE. These findings provide a strong preclinical rationale for clinical development of [^225^Ac]Ac-SSO110 in SSTR2-positive malignancies.

## DISCLOSURE

This study was funded by Ariceum Therapeutics GmbH. The authors listed in the study are employees of companies within the Ariceum group. Ariceum Therapeutics GmbH holds patent applications related to the technology described in this work (including WO2024121249A1 and WO2025255396A1), on which some of the authors are listed as inventors.

## ACKNOWLEDGMENTS

The authors thank Advancell, Oncodesign Services and Chelatec for conducting the in vivo studies. The authors are grateful to Nicolas Chouin for support with tumor-absorbed dose calculations. Bilal Bham is acknowledged for contributions in manuscript preparation, and Shelby Hartzell for language editing and review.

## REFERENCES

1. Pauwels E, Cleeren F, Bormans G, Deroose CM. Somatostatin receptor PET ligands - the next generation for clinical practice. Am J Nucl Med Mol Imaging. 2018;8:311–331.

2. Hirmas N, Jadaan R, Al-Ibraheem A. Peptide Receptor Radionuclide Therapy and the Treatment of Gastroentero-pancreatic Neuroendocrine Tumors: Current Findings and Future Perspectives. Nucl Med Mol Imaging. 2018;52:190–199.

3. Watanabe N, Nakanishi Y, Kinukawa N, et al. Expressions of Somatostatin Receptor Subtypes (SSTR-1, 2, 3, 4 and 5) in Neuroblastic Tumors; Special Reference to Clinicopathological Correlations with International Neuroblastoma Pathology Classification and Outcomes. Acta Histochem Cytochem. 2014;47:219–229.

4. Remes SM, Leijon HL, Vesterinen TJ, Arola JT, Haglund CH. Immunohistochemical Expression of Somatostatin Receptor Subtypes in a Panel of Neuroendocrine Neoplasias. J Histochem Cytochem. 2019;67:735–743.

5. Priyadarshini S, Allison DB, Chauhan A. Comprehensive Assessment of Somatostatin Receptors in Various Neoplasms: A Systematic Review. Pharmaceutics. 2022;14:1394.

6. Han G, Hwang E, Lin F, et al. RYZ101 (Ac-225 DOTATATE) Opportunity beyond Gastroenteropancreatic Neuroendocrine Tumors: Preclinical Efficacy in Small-Cell Lung Cancer. Molecular Cancer Therapeutics. 2023;22:1434–1443.

7. Fagerstedt KW, Vesterinen T, Leijon H, Sihto H, Böhling T, Arola J. Somatostatin receptor expression in Merkel cell carcinoma: correlation with clinical data. Acta Oncologica. 2023;62:1001–1007.

8. Kersting D, Sandach P, Sraieb M, et al. 68Ga-SSO-120 PET for Initial Staging of Small Cell Lung Cancer Patients: A Single-Center Retrospective Study. J Nucl Med. 2023;64:1540–1549.

9. Mavroeidi IA, Romanowicz A, Haake T, et al. Theranostics with somatostatin receptor antagonists in SCLC: Correlation of 68Ga-SSO120 PET with immunohistochemistry and survival. Theranostics. 2024;14:5400–5412.

10. Strosberg JR, Caplin ME, Kunz PL, et al. 177Lu-Dotatate plus long-acting octreotide versus high-dose long-acting octreotide in patients with midgut neuroendocrine tumours (NETTER-1): final overall survival and long-term safety results from an open-label, randomised, controlled, phase 3 trial. Lancet Oncol. 2021;22:1752–1763.

11. Strosberg J, El-Haddad G, Wolin E, et al. Phase 3 Trial of^177^ Lu-Dotatate for Midgut Neuroendocrine Tumors. N Engl J Med. 2017;376:125–135.

12. Ginj M, Zhang H, Waser B, et al. Radiolabeled somatostatin receptor antagonists are preferable to agonists for *in vivo* peptide receptor targeting of tumors. Proc Natl Acad Sci USA. 2006;103:16436–16441.

13. Dalm SU, Nonnekens J, Doeswijk GN, et al. Comparison of the Therapeutic Response to Treatment with a^177^ Lu-Labeled Somatostatin Receptor Agonist and Antagonist in Preclinical Models. J Nucl Med. 2016;57:260–265.

14. Nicolas GP, Mansi R, McDougall L, et al. Biodistribution, Pharmacokinetics, and Dosimetry of^177^ Lu-,^90^ Y-, and^111^ In-Labeled Somatostatin Receptor Antagonist OPS201 in Comparison to the Agonist^177^ Lu-DOTATATE: The Mass Effect. J Nucl Med. 2017;58:1435–1441.

15. Albrecht J, Exner S, Grötzinger C, et al. Multimodal Imaging of 2-Cycle PRRT with^177^ Lu-DOTA-JR11 and^177^ Lu-DOTATOC in an Orthotopic Neuroendocrine Xenograft Tumor Mouse Model. J Nucl Med. 2021;62:393–398.

16. Mansi R, Plas P, Vauquelin G, Fani M. Distinct In Vitro Binding Profile of the Somatostatin Receptor Subtype 2 Antagonist [177Lu]Lu-OPS201 Compared to the Agonist [177Lu]Lu-DOTA-TATE. Pharmaceuticals. 2021;14:1265.

17. Plas P, Limana L, Carré D, et al. Comparison of the Anti-Tumour Activity of the Somatostatin Receptor (SST) Antagonist [177Lu]Lu-Satoreotide Tetraxetan and the Agonist [177Lu]Lu-DOTA-TATE in Mice Bearing AR42J SST2-Positive Tumours. Pharmaceuticals. 2022;15:1085.

18. Wild D, Fani M, Fischer R, et al. Comparison of Somatostatin Receptor Agonist and Antagonist for Peptide Receptor Radionuclide Therapy: A Pilot Study. J Nucl Med. 2014;55:1248–1252.

19. Tafreshi NK, Doligalski ML, Tichacek CJ, et al. Development of Targeted Alpha Particle Therapy for Solid Tumors. Molecules. 2019;24:4314.

20. Pouget J-P, Constanzo J. Revisiting the Radiobiology of Targeted Alpha Therapy. Front Med (Lausanne). 2021;8:692436.

21. Borgna F, Barritt P, Grundler PV, et al. Simultaneous Visualization of 161Tb- and 177Lu-Labeled Somatostatin Analogues Using Dual-Isotope SPECT Imaging. Pharmaceutics. 2021;13:536.

22. Busslinger SD, Mapanao AK, Kegler K, et al. Comparison of the tolerability of 161Tb-and 177Lu-labeled somatostatin analogues in the preclinical setting. Eur J Nucl Med Mol Imaging. July 2024.

23. Lehenberger S, Barkhausen C, Cohrs S, et al. The low-energy β− and electron emitter 161Tb as an alternative to 177Lu for targeted radionuclide therapy. Nuclear Medicine and Biology. 2011;38:917–924.

24. Alcocer-Ávila ME, Ferreira A, Quinto MA, Morgat C, Hindié E, Champion C. Radiation doses from 161Tb and 177Lu in single tumour cells and micrometastases. EJNMMI Phys. 2020;7:33.

25. Borgna F, Haller S, Rodriguez JMM, et al. Combination of terbium-161 with somatostatin receptor antagonists—a potential paradigm shift for the treatment of neuroendocrine neoplasms. Eur J Nucl Med Mol Imaging. 2022;49:1113–1126.

26. Song Z, Zhang J, Qin S, et al. Targeted alpha therapy: a comprehensive analysis of the biological effects from “local-regional-systemic” dimensions. Eur J Nucl Med Mol Imaging. 2025;53:30–47.

27. Stallons TAR, Saidi A, Tworowska I, Delpassand ES, Torgue JJ. Preclinical Investigation of 212Pb-DOTAMTATE for Peptide Receptor Radionuclide Therapy in a Neuroendocrine Tumor Model. Mol Cancer Ther. 2019;18:1012–1021.

28. Handula M, Beekman S, Konijnenberg M, et al. First preclinical evaluation of [225Ac]Ac-DOTA-JR11 and comparison with [177Lu]Lu-DOTA-JR11, alpha versus beta radionuclide therapy of NETs. EJNMMI Radiopharm Chem. 2023;8:13.

29. Wild D, Grønbæk H, Navalkissoor S, et al. A phase I/II study of the safety and efficacy of [177Lu]Lu-satoreotide tetraxetan in advanced somatostatin receptor-positive neuroendocrine tumours. Eur J Nucl Med Mol Imaging. 2023;51:183–195.

