## Supplemental Data for "[^225^Ac]Ac-SSO110 demonstrates superior efficacy over ^177^Lu-, ^161^Tb-, and ^212^Pb-labeled SSO110 and [^225^Ac]Ac-DOTA-TATE in SSTR2-positive tumor models"

**Supplemental methods**

**Radiochemistry**

***[$^{212}$Pb]Pb-SSO110:*** In an Eppendorf tube, [$^{212}$Pb]$PbCl_3$ (Advancell, 0.05 M HCl) was mixed with radiolabeling buffer (pH 5.5) containing SSO110 and heated at 95 °C for 25 minutes. The reaction mixture was cooled to room temperature, and a stabilizing solution with 100 mg/mL sodium ascorbate and 0.1 mg/mL DTPA was added. The final specific activity ranged from 0.1 to 0.75 MBq/µg.

***$^{161}$Tb-, and $^{177}$Lu-SSO110***: In a microtube, [$^{161}$Tb]$TbCl_3$ (Terthera, 0.05 M HCl) or [$^{177}$Lu]$LuCl_3$ (ITM/Eckert & Ziegler, 0.04 M HCl) was mixed with radiolabeling buffer (pH 5.1) and SSO110 (dissolved in 0.1% TFA). The reaction mixture was incubated at 90 °C for 30 minutes. At the end of the incubation, 4 mM DTPA in 0.9% NaCl was added at room temperature to chelate unbound $^{161}$Tb or $^{177}$Lu. Final solutions were diluted in 0.9% NaCl supplemented with 5 mg/mL ascorbic acid. The final specific activity ranged from 10 to 20 MBq/µg for [$^{161}$Tb]Tb-SSO110 and 20 MBq/µg for [$^{177}$Lu]Lu-SSO110.

***[$^{225}$Ac]Ac-SSO110 and [$^{225}$Ac]Ac-DOTA-TATE:*** In an Eppendorf tube, [$^{225}$Ac]$AcCl_3$ (ITM, 0.04 M HCl) was mixed with radiolabeling buffer (pH 8.1) and SSO110 or DOTA-TATE and heated at 80 °C for 20-30 minutes. The reaction mixture was cooled to room temperature, followed by addition of a stabilizing solution containing 100 mg/mL sodium ascorbate and 0.1 mg/mL DTPA. The final specific activity ranged from 3.3 to 90 kBq/µg for [$^{225}$Ac]Ac-SSO110 and from 21 to 42 kBq/µg for [$^{225}$Ac]Ac-DOTA-TATE.

Radiolabeling incorporation and radiochemical purity was assessed by iTLC and HPLC, respectively, for all radioconjugates.

**Cell lines and tumor models**

For in vivo biodistribution and efficacy studies, female Balb/c nude or Swiss nude mice (6–8 weeks old; Janvier Labs or Charles River) were used. NCI-H69 and AR42J cells (ECACC) were cultured in RPMI 1640 supplemented with 10% fetal bovine serum (AR42J additionally with 2 mM L-glutamine) and were confirmed to be mycoplasma-free prior to implantation.

**In vivo studies**

For xenograft studies, cells were harvested during exponential growth phase using standard tissue culture procedures and were confirmed to be mycoplasma-free prior to implantation. Balb/c nude mice were subcutaneously implanted in the right flank with $5 \times 10^6$ NCI-H69 cells suspended in PBS/Matrigel (1:1). Swiss nude mice received $1 \times 10^7$ AR42J cells in phenol red-free RPMI 1640 medium. AR42J implantation was performed 48 h after whole-body irradiation (2 Gy, $^{60}$Co). Animals were randomized one day prior to treatment.

Tumor volumes (mm$^3$) were measured at baseline and at least twice weekly thereafter using calipers and calculated using the formula (width$^2$ × length)*0.5. Median survival was assessed by Kaplan-Meier analysis, with death recorded as event = 1 and surviving animals censored as event = 0. Body weight was monitored individually on the days indicated in the graphs. Animals were euthanized when tumor volumes reached 1500 mm$^3$ (AR42J xenografts) or 1000 mm$^3$ (NCI-H69 xenografts), or earlier in the event of tumor ulceration or sustained body weight loss ≥ 20% for two consecutive days.

For efficacy studies, blood samples were collected at study termination or upon reaching humane endpoints and processed to obtain serum (AR42J model) or plasma (NCI-H69 model) for analysis of albumin (ALB), creatinine (CREA), blood urea nitrogen (BUN), alkaline phosphatase (ALP), and total bilirubin (TBIL). Values below the lower limit of detection (LLOD) were assigned to a value of LLOD*0.5.

For histopathological evaluation, paraffin-embedded tumor sections (4 µm thickness) from AR42J-bearing mice treated with vehicle, 37 kBq [$^{225}$Ac]Ac-SSO110, or 37 kBq [$^{225}$Ac]Ac-DOTA-TATE were stained with hematoxylin and eosin. Histopathological analysis was performed by a veterinary pathologist using a predefined scoring system.

All animal experiments were approved by the relevant institutional and national authorities for animal welfare and were conducted in accordance with applicable guidelines.

**Tumor absorbed dose calculations**

The mean absorbed doses of [$^{225}$Ac]Ac-SSO110 and [$^{212}$Pb]Pb-SSO110 to the tumor was calculated using the MIRD formalism (*1*). Time–activity curves from biodistribution data were fitted with either a mono-exponential function or a bi-exponential using GraphPad PRISM (GraphPad Software Inc., San Diego, CA, USA), with fitting selection based on Akaike's Informative Criteria. The time-integrated activity in the tumor was calculated by integrating the selected function from time zero to infinity. Only the contribution of α-particles to the tumor-absorbed dose was considered, as β-particle contributions were assumed to be negligible. Therefore, crossfire effects between tissues were not accounted for. The tumor absorbed doses were calculated by multiplying the time-integrated activity per gram of tumor (Bq*s/g) by the total α-particle energy emitted per decay for $^{225}$Ac (27.5 MeV) and $^{212}$Pb (7.8 MeV).

**Immunohistochemistry of tissue samples**

Xenograft tumor tissues (NCI-H69 and AR42J) collected from mice enrolled in efficacy studies were processed into formalin-fixed paraffin-embedded (FFPE) tissue sections according to standard procedures. Anti-SSTR2 immunohistochemistry (IHC) was performed using the Discovery Ultra staining platform. Briefly, following deparaffinization and heat-induced antigen retrieval with citrate buffer, pH 6 for 32 minutes at 91 °C, sections were incubated with rabbit anti-SSTR2 antibody (clone UMB1, 6 μg/mL) for 60 minutes at 35 °C. Sections were then incubated with OmniMap anti-rabbit HRP secondary antibody for 16 minutes, followed by DAB chromogenic detection. Nuclei were counterstained with Hematoxylin II for 8 minutes and exposed to Bluing reagent for 4 minutes. Rabbit IgG (clone DA1E, 6 μg/mL) was used as an isotype control. SSTR2 expression was semi-quantitatively evaluated using H-Score analysis by estimating the percentages of weak, moderate, and strong staining in tumor and non-malignant cells. The H-score in the range from 0 to 300 was calculated as:

$$H - score\ = (\%\, weak)\ + 2\ \times (\%\, moderate)\ + 3\ \times (\%\, strong)$$

**Statistical analysis**

Statistical analyses were performed using GraphPad Prism software. Quantitative histopathologic data was analyzed using ordinary one-way ANOVA followed by Tukey's multiple comparison test. Survival curves were generated using the Kaplan-Meier method, and overall differences between groups was assessed using the log-rank test. Clinical chemistry parameters were compared between treatment groups and their respective vehicle controls using one-way ANOVA followed by Dunnett's multiple comparison test. Statistical significance is either represented by p values or asterisks. *, **, ***, **** represents $p \leq 0.05$, 0.01, 0.001, and 0.0001, respectively. Values of $p > 0.05$ were considered not statistically significant.

### Supplemental figures

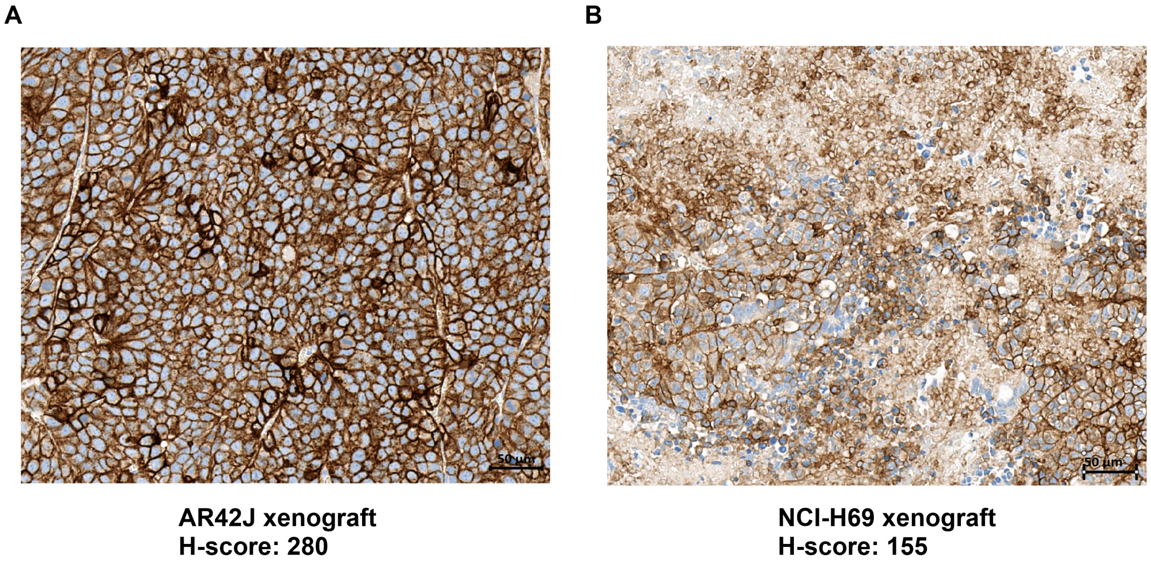


**Figure S1.** Representative immunohistochemical staining of SSTR2 in tumor xenografts. Tumor sections from (A) AR42J and (B) NCI-H69 xenografts were stained with an anti-SSTR2 antibody. Corresponding H-scores of the representative images are shown. Scale bar: 50 µm

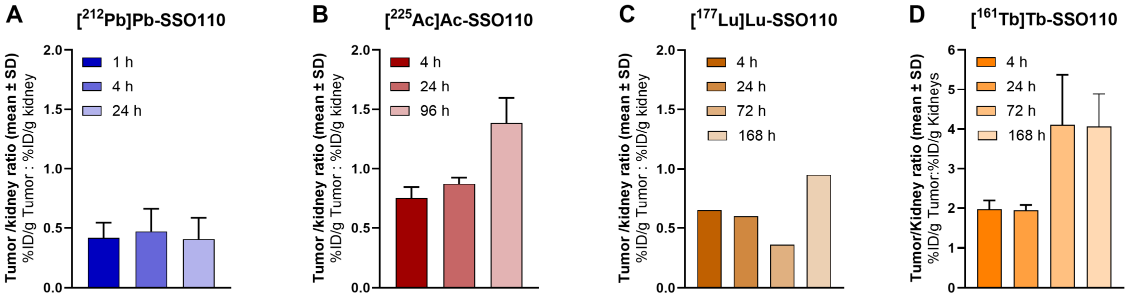


**Figure S2**. Tumor-to-kidney ratios over time. Ratios were determined in mice treated with (A) [$^{212}$Pb]Pb-SSO110, (B) [$^{225}$Ac]Ac-SSO110, (C) [$^{177}$Lu]Lu-SSO110, and (D) [$^{161}$Tb]Tb-SSO110. Data are presented as mean ± SD.

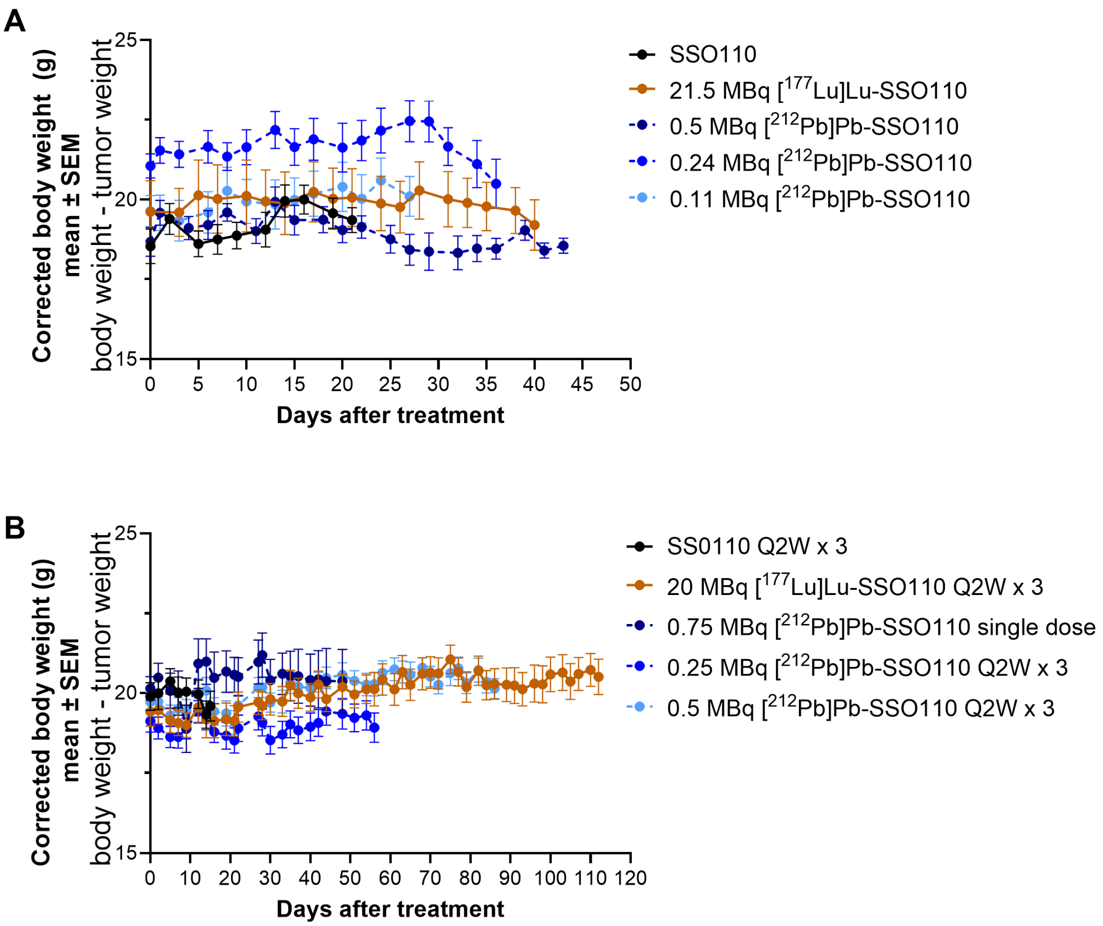


**Figure S3**. Body weight changes in the NCI-H69 model following treatment. NCI-H69 tumor-bearing mice were treated with single doses of (A) non-radiolabeled SSO110, [$^{177}$Lu]Lu-SSO110, or [$^{212}$Pb]Pb-SSO110. B) NCI-H69-bearing mice were treated with multiple doses (Q2W x 3) of SSO110, [$^{177}$Lu]Lu-SSO110, or [$^{212}$Pb]Pb-SSO110. Body weights were corrected for tumor volume and plotted until the first animal dropout. Data are presented as mean ± SEM.

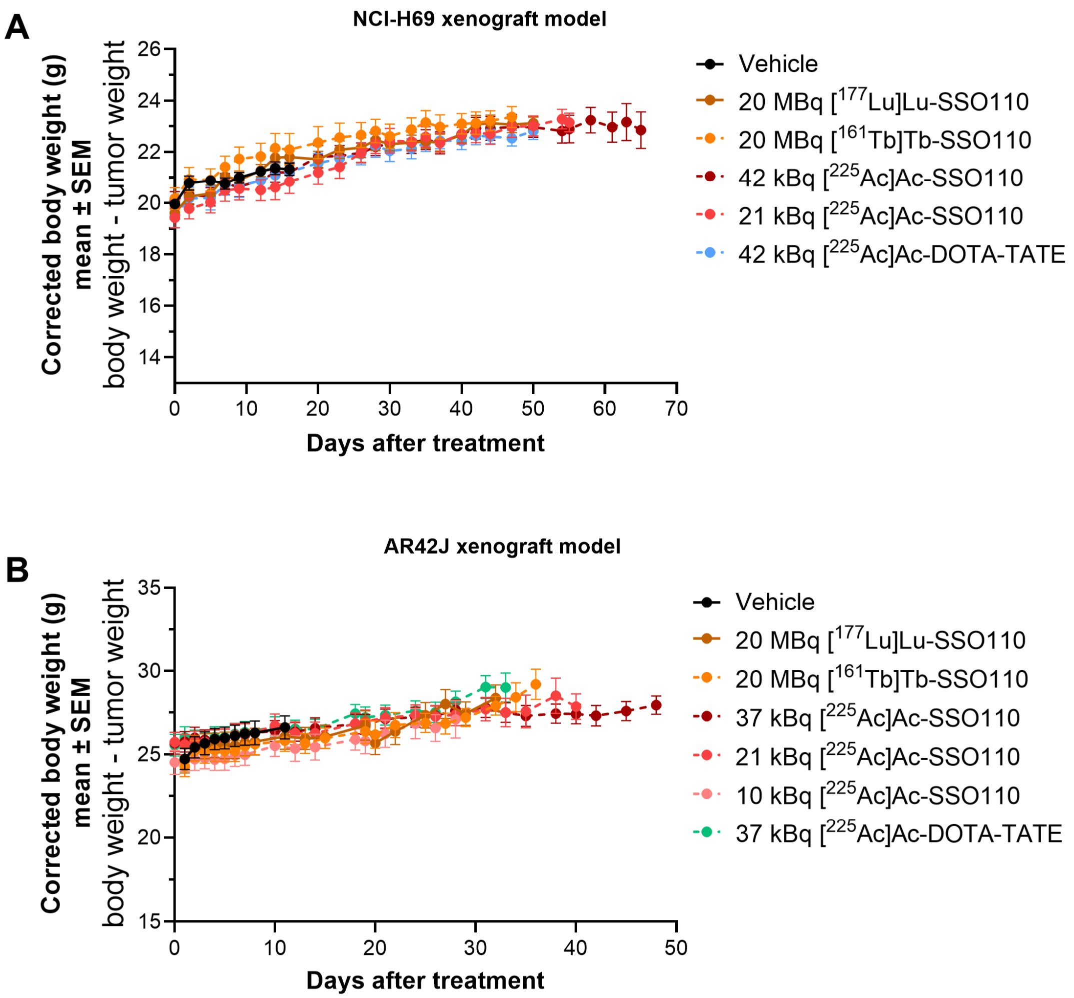


**Figure S4**. Body weight changes in the NCI-H69 and AR42J model following treatment. (A) NCI-H69 tumor-bearing mice were treated with vehicle, $^{177}$Lu-, $^{161}$Tb-, and $^{225}$Ac-labeled SSO110, or [$^{225}$Ac]Ac-DOTA-TATE. (B) AR42J tumor-bearing mice were treated with vehicle, $^{177}$Lu-, $^{161}$Tb-, and $^{225}$Ac-labeled SSO110, or [$^{225}$Ac]Ac-DOTA-TATE. Body weights were corrected for tumor volume and plotted until the first animal dropout (NCI-H69) or until 50% of animals remained in the study (AR42J). Data are presented as mean ± SEM.

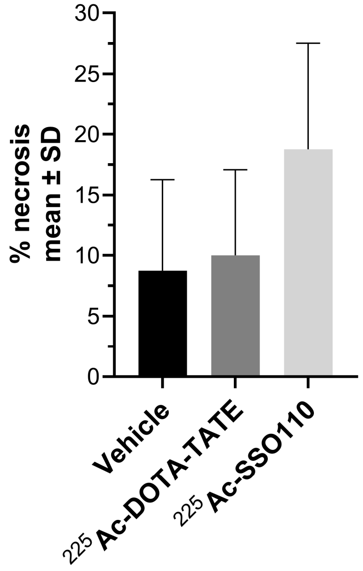


**Figure S5**. Histopathological analysis of AR42J xenograft tumors after treatment with vehicle, 37 kBq [$^{225}$Ac]Ac-DOTA-TATE or [$^{225}$Ac]Ac-SSO110. Shown is the percentage of necrotic neoplastic cells. Data are presented as mean ± SD.

1. Bolch WE, Eckerman KF, Sgouros G, Thomas SR. MIRD pamphlet No. 21: a generalized schema for radiopharmaceutical dosimetry--standardization of nomenclature. *J Nucl Med*. 2009;50:477-484.
